# Serum GlycA level is a candidate biomarker for disease activity in systemic lupus erythematosus and for proliferative status of lupus nephritis, independent of renal function impairment

**DOI:** 10.1101/493809

**Authors:** Tim Dierckx, Sylvie Goletti, Laurent Chiche, Laurent Daniel, Bernard Lauwerys, Noémie Jourde-Chiche, Johan Van Weyenbergh

## Abstract

**Objective:** Glycoprotein acetylation (GlycA) is a novel biomarker for chronic inflammation, associated to cardiovascular risk. Serum GlycA levels are increased in several inflammatory diseases, including systemic lupus erythematosus (SLE). We investigated the relevance of serum GlycA measurement in SLE and lupus nephritis (LN).

**Methods:** GlycA was measured by NMR in 194 serum samples from patients and controls. Comparisons were performed between groups. Clinical and biological parameters were tested for correlation with GlycA levels. The predictive value of GlycA to differentiate proliferative from non-proliferative LN was determined using logistic regression models.

**Results:** GlycA was correlated to C-reactive protein (CRP), neutrophil count, proteinuria and the SLE disease activity index (SLEDAI), and inversely with serum albumin. GlycA was higher in active (n=105) than in quiescent (n=39) SLE patients, in healthy controls (n=29), and in patients with non-lupus nephritis (n=21), despite a more altered renal function in the latter. In patients with biopsy-proven active LN, GlycA was higher in proliferative (n=32) than non-proliferative (n=11) LN, independent of renal function and proteinuria level. Logistic regression models showed that, in univariate models, GlycA outperforms traditional biomarkers. A bivariate model using GlycA and BMI better predicted the proliferative status of LN than a model comprising CRP, renal function (eGFR), serum albumin, proteinuria, C3 consumption and the presence of anti-dsDNA antibodies.

**Conclusion:** Serum GlycA is elevated in SLE, and correlates with disease activity and LN. Serum GlycA, which summarizes different inflammatory processes, could be a valuable biomarker to discriminate proliferative from non-proliferative LN and should be tested in large, prospective cohorts.

## Introduction

Systemic inflammation is implicated in a wide range of auto-immune diseases such as systemic lupus erythematosus (SLE) [1]. In SLE, a multitude of different inflammatory cytokines and pathways can be activated [2] and these are not always reflected by elevated C-reactive protein (CRP). Lupus nephritis (LN) is one of the most severe complications of SLE which occurs in 20-70% of patients and its occurrence is not reliably predicted by classical inflammatory markers [1]. The occurrence and severity of LN has been associated with the dysregulation of neutrophil extra-cellular traps (NETs) [3] and a blood transcriptional neutrophil signature [4]. Non-invasive biomarkers are needed in SLE to predict the risk of LN in SLE patients, and to determine the severity of LN by differentiating between the proliferative and non-proliferative forms of LN, which determines whether immunosuppressive therapy will be needed to achieve renal remission [5].

Glycoprotein Acetylation (GlycA) is a new nuclear magnetic resonance (NMR) spectroscopy-derived biomarker of systemic inflammation that reflects protein glycosylation which could prove to be promising biomarker in the LN context. GlycA signal, measured in the blood serum or plasma, reflects mainly the glycosylation of acute-phase proteins α1-acid glycoprotein, haptoglobin, α1antitrypsin, α1antichymotrypsin and transferrin [6], [7], as a consequence of inflammatory stimuli. The GlycA signal is thus evaluated as a biomarker of systemic inflammation and cardiovascular risk [8], summarizing the activation of multiple inflammatory pathways [9]. GlycA has been shown to be associated with the risk of cardio-vascular events and mortality in the general population, independently of CRP [10], [11]. In a large Brazilian cohort, GlycA was associated with CRP, age, female gender, tobacco and alcohol consumption, obesity, diabetes, hypertension and dyslipidemia and associated independently with lower estimated glomerular filtration rate (eGFR) and albuminuria [12]. We recently showed that GlycA could be a marker of disease activity in inflammatory bowel disease, even in patients without CRP elevation [13]. Of particular interest to the LN context, large scale gene correlation network analysis has shown that GlycA can also be associated to NET formation [14]. Furthermore, concentrations of GlycA were noted to be higher in SLE patients than in healthy controls, but findings with regards to GlycA’s association to disease activity vary between studies [15], [16]. The value of GlycA as a biomarker of LN has not been evaluated yet.

In this work, we investigated if GlycA could be associated with LN severity. We compared patients with proliferative and non-proliferative forms of biopsy-proven LN, and included a control group of patients with non-lupus renal diseases to account for the possible confounding factors of altered eGFR and albuminuria.

## Materials and Methods

### Patient demographics & Ethics

This study was conducted in accordance with the principles of the declaration of Helsinki.

Patients with biopsy-proven LN, and patients with other renal diseases, were included in the biobank DC-2012-1704, approved by the French Ministry of Health, in the Hôpital de la Conception, Marseille, France. All patients gave written informed consent before any procedure.

Other samples from patients with SLE were obtained from patients recruited in the LOUvain Lupus Nephritis InCeption (LOULUNIC) cohort, and patients followed-up at the Lupus Clinic of the Université Catholique de Louvain (UCL), Brussels, Belgium, as were samples from age- and sex-matched healthy controls. All patients and controls gave written informed consent before serum samplings. All patients with SLE responded to the SLICC 2012 classification criteria [17].

### Clinical metrics

For this study, a total of n=194 samples were analyzed. A graphical overview of assayed samples can be found in **Figure 1**. We grouped the samples as either originating from healthy controls, from patients with biopsy-proven non-lupus renal diseases (‘non-lupus nephritic controls’, comprising patients with membranous nephropathy, IgA nephropathy, diabetic kidney disease or hypertensive nephropathy) and from SLE patients. SLE patients were further identified as clinically quiescent SLE patients (SLEDAI =< 4 without clinical activity, with or without maintenance therapy, immunological activity authorized), or active SLE patients. Active SLE patients were either SLE patients with an extra-renal flare without nephritic involvement, or patients with active LN (whether there is extra-renal activity or not). For differential analysis between proliferative (class III or IV +/− V, with active lesions, of the ISN/RPS 2003 classification) and non-proliferative (class I, II or isolated class V) flaring LN cases, a subset of samples was used, which originates from patients who were sampled at the time of the biopsy-proven LN. Patient numbers for this cohort, as well as full demographical and clinical details, can be found in **Supplementary Tables 1** & 2. For 28 SLE patients, longitudinal samples were available.

**Figure 1:**
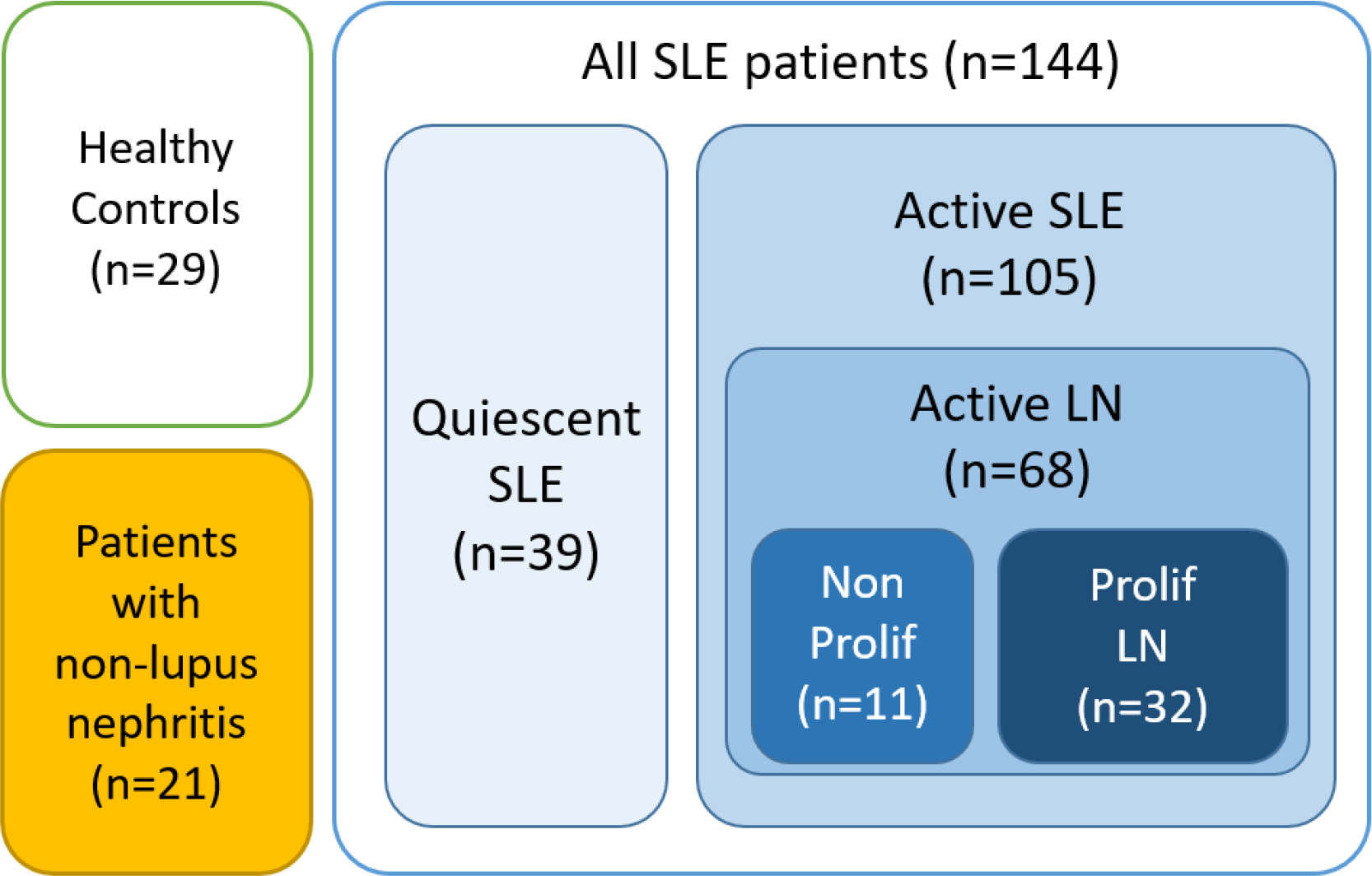
Sample overview.

Age, Gender, ethnicity, BMI, smoking status, extra-renal lupus activity (ongoing arthritic flare in particular), and diabetic status, were collected and tested for confounding effects on GlycA. Serum CRP level, serum albumin and creatinine levels, estimated glomerular filtration rate (eGFR, calculated with the MDRD equation [18], C3 and C4 concentration, urinary protein/creatinine ratio (UPCR), presence of anti-dsDNA antibodies as a binary variable (above 16 UI/mL by ELIA^™^, ThermoFisher, MA, USA, or above 10 IU/mL using the Farr assay from Trinity Biotech, Bray, Ireland), daily glucocorticoid dosage, and hydroxychloroquine usage were tested for association to GlycA on all available SLE samples.

GlycA concentration was quantified using the Nightingale Health Ltd. high-throughput metabolomics platform (Helsinki, Finland) [6], [7]. Laboratory and NMR measurements of creatinine and albumin concentrations were found to be highly correlated (*ρ*=0.94 and *ρ*=0.74, respectively, with p-values<10^−8^) (**Supplementary Figure 1**).

### Statistical analysis

Univariate comparison between the different conditions was performed using unpaired, two-sided Wilcoxon rank sum test. Association between GlycA and all available clinical and demographic data was tested using Spearman’s *ρ* for continuous variables and Mann-Whitney-Wilcoxon’s U for categorical variables, p values were corrected for multiple testing using the Benjamini-Hochberg procedure and resulting q values are accepted as significant when smaller than 0.05. Identification of all confounding factors on the relationship between GlycA and proliferative status was achieved with analysis of covariance methods calculating the Type III Sums of Squares in a stepwise forward selection. Starting from the model including only the GlycA response variable and proliferative status, in each step the variable with the most significant contribution (with maximum *p*-value 0.1), as determined in a comparison with an F-test, was included in the model. Discriminatory power of clinical and demographical parameters was assessed in logit-link logistic regression models of LN patients with proliferative status of the patient as the response variable. In addition to univariate and selected multivariate models, a statistically ideal logistic regression model was constructed using an exhaustive best subset algorithm using Aikake’s Information Criterion [19]. All statistical analyses were performed in R [20]. C statistics for logistic regression models were calculated using the Epi R package [21]. Leave-one-out cross-validated accuracy measures were calculated using the caret R package [22]. Figures were generated using the ggplot2 R package [23].

## Results

### GlycA levels in SLE patients and controls

Patients with active SLE showed significantly higher GlycA concentration than healthy controls (p=0.009), non-lupus nephritic controls (p=0.04) and quiescent SLE patients (p<10^−6^) (**Figure 2**). Quiescent SLE patients also displayed lower CRP levels than patients with active SLE (p<10^−3^). However, in contrast to GlycA levels, CRP concentrations of non-lupus nephritic controls were not significantly different from those of active SLE patients (p=0.86). eGFR was significantly lower in non-lupus nephritic controls than in active SLE patients (p<10^−3^), indicating that the increased GlycA levels observed in active SLE can’t be solely attributed to a decrease in renal function (**Figure 2**).

**Figure 2:**
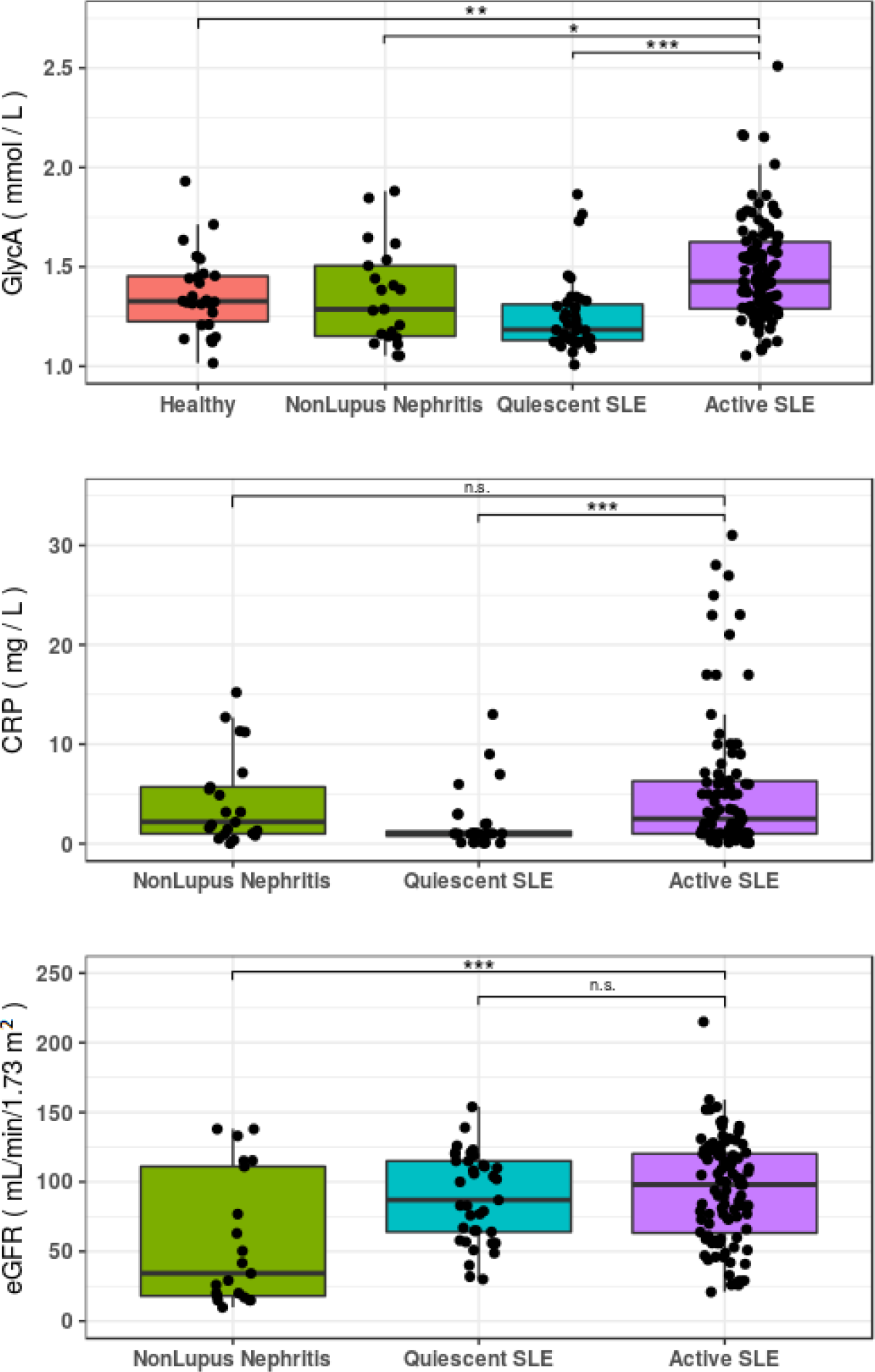
GlycA is elevated in active SLE, compared to healthy controls, non-lupus nephritic controls, and quiescent SLE. C-reactive protein (CRP) level is lower in quiescent SLE than in active SLE and non-lupus nephritis but does not differ between active SLE and non-lupus nephritis. Estimated glomerular filtration rate (eGFR) is lower in patients with non-lupus nephritis than in patients with quiescent or active SLE, excluding the fact that the elevation of GlycA in active SLE may be explained solely by decreased renal function. Significance of a Wilcoxon test comparing the groups to active SLE are indicated (* p < 0.05, ** p < 0.01, *** p < 0.001).

### GlycA association to clinical measurements

GlycA correlated well with CRP in all samples (ρ=0.49, q<10^−8^). Significant correlations to GlycA were observed for neutrophil counts (ρ=0.32, q=0.04), serum albumin (ρ=−0.37, q<10^−4^), and UPCR (ρ=0.38, q<10^−4^), and trends for correlation were observed for serum creatinine (ρ=0.17, q=0.06) and C3 (ρ=−0.18, q=0.06). GlycA showed no correlation to anti-dsDNA levels measured by ELIA^™^ or Farr, and no significant difference was found between GlycA concentrations of dsDNA positive SLE patients, when compared to dsDNA negative SLE patients. GlycA levels showed a trend for inverse correlation with eGFR in quiescent SLE patients (ρ=−0.37, q=0.07) and in non-lupus nephritic controls with a wide range of eGFR and proteinuria levels (ρ=−0.46, q=0.06). In SLE samples, GlycA showed a significant correlation to the SLEDAI score (ρ=0.36, q<10^−4^). This correlation between the SLEDAI score and GlycA was absent if only quiescent SLE samples were considered. The full results of correlational analysis with GlycA can be found in **Supplementary Table 3**. **Figure 3** shows GlycA levels of available longitudinal samples of LN patients with a flare event. While insufficient observations are available to perform robust longitudinal analysis, we note that GlycA level is variable over time in a single patient, and that patients show increased GlycA at or directly following the flare presentation. Most patients show a return to pre-flare GlycA levels following the flare resolution.

**Figure 3:**
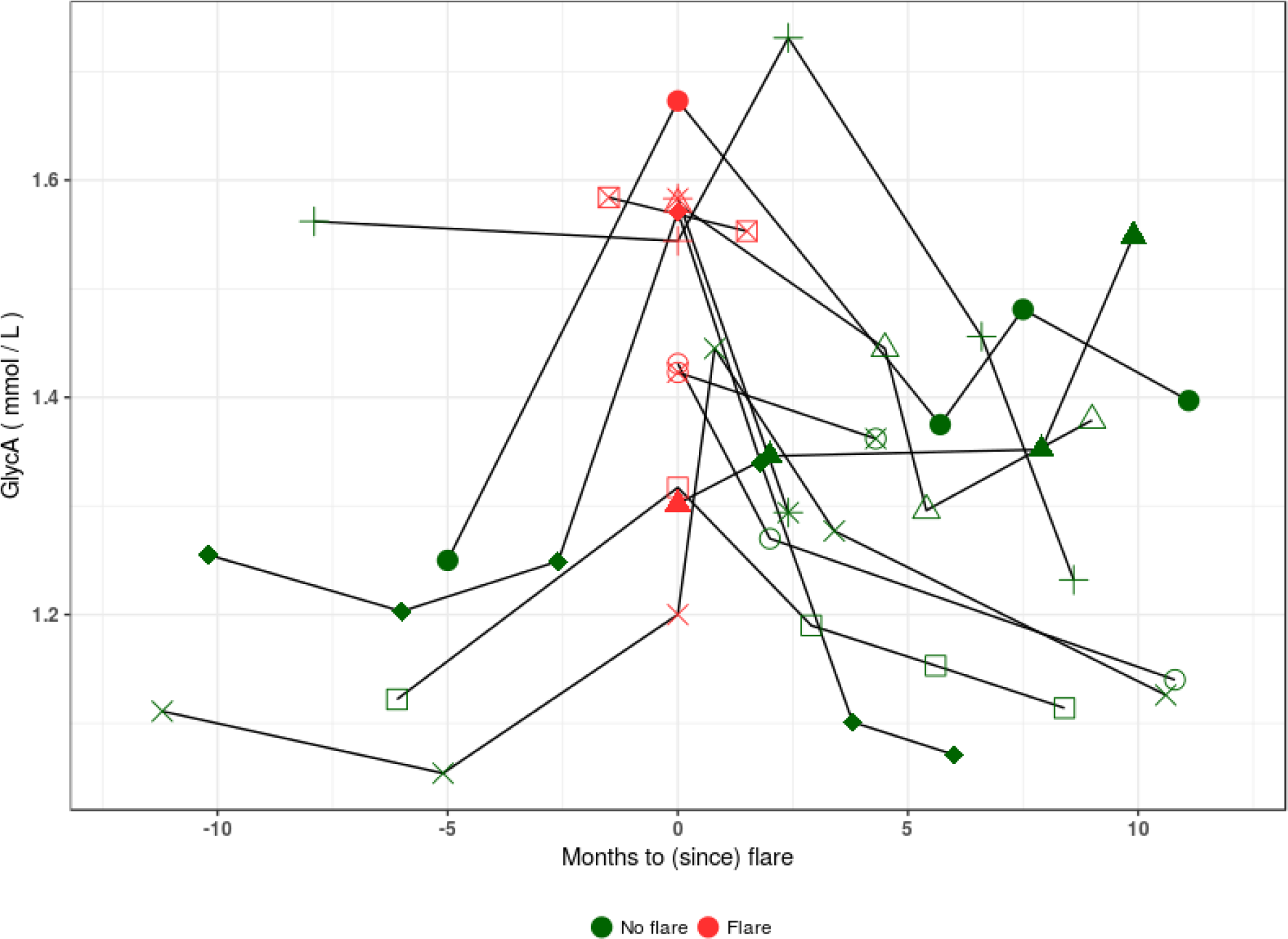
GlycA levels measured in longitudinal samples of flaring LN patients, up to a year prior and after flare event. Patients show increased GlycA levels at or directly following flare presentation. Post flare, most patients show a return to maintenance GlycA levels.

### GlycA association to proliferative status in flaring LN

GlycA concentrations in serum samples taken at time of a flare of patients with biopsy-proven active LN were higher in proliferative cases than in non-proliferative cases (p=0.04). Although flaring proliferative LN cases had lower eGFR than non-proliferative cases, this difference was not statistically significant (p=0.28) and non-lupus nephritic controls with considerably lower eGFR than the proliferative LN patients (p<0.01) had GlycA levels comparable to those of healthy controls and non-proliferative flaring LN (**Figure 4**).

**Figure 4:**
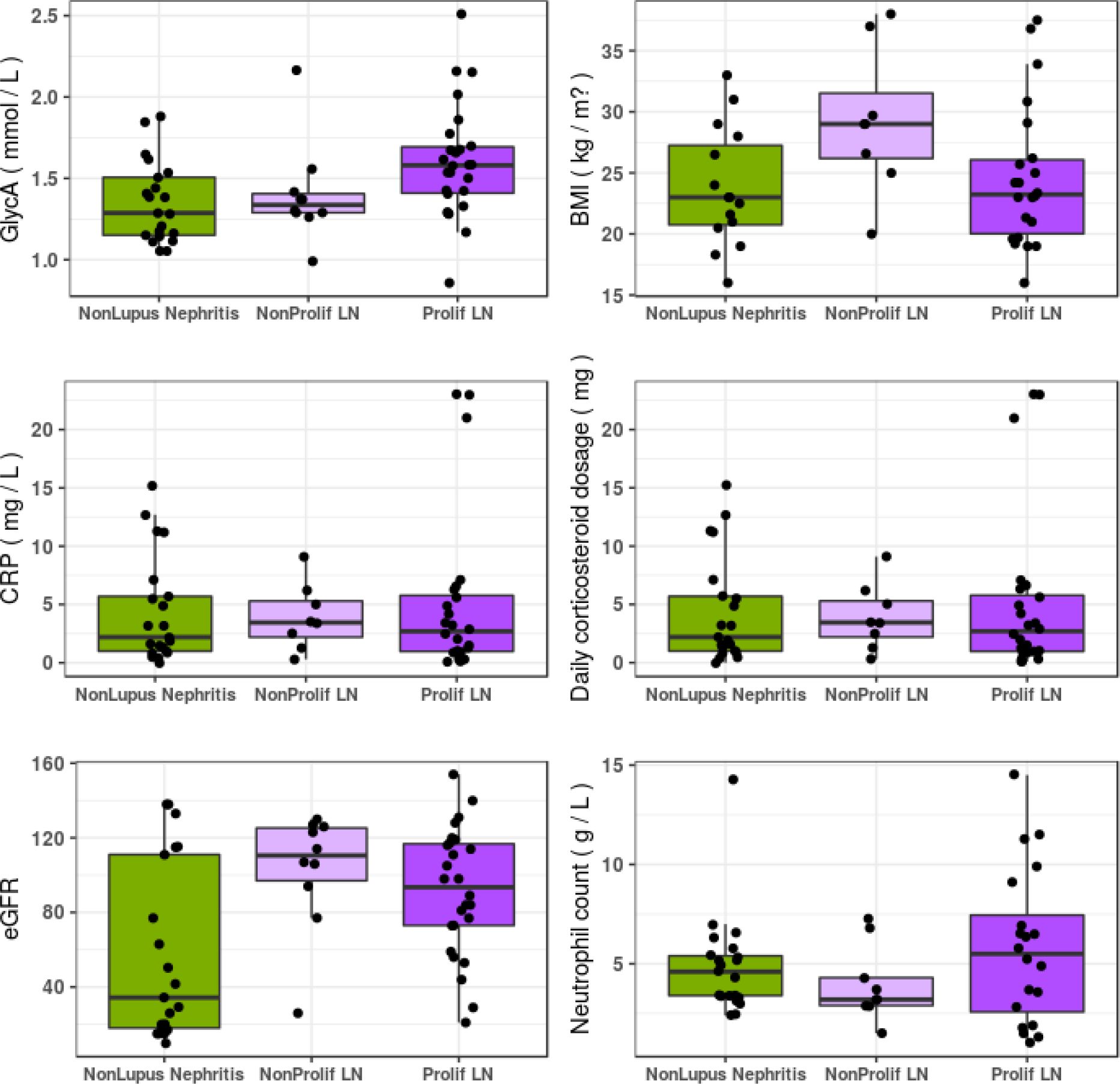
Comparison between LN samples taken at flare with biopsy-proven proliferative (or non-proliferative) status within 30 days of sampling shows significantly increased GlycA in proliferative LN. In addition, CRP and eGFR, as well as identified confounding factors BMI, daily corticosteroid dosage and neutrophil count are visualized in the flaring samples as well as in non-lupus nephritic control samples.

We sought to identify possible confounding factors interfering with GlycA levels’ association to proliferative status of patients with flaring LN, such as BMI, which is known to be associated to GlycA, and was significantly higher in patients with non-proliferative than proliferative LN in this cohort (p=0.04). Analysis of variance identified confounding effects of BMI, daily corticosteroid dosage and neutrophil count as having independent and significant impacts on proliferative status’ association to GlycA levels. After controlling for BMI, daily corticosteroid dosage and neutrophil concentration, proliferative status retained its association to GlycA.

### Logistic regression models of proliferative status

We assessed the discriminatory power of GlycA in logistic regression models built to differentiate between proliferative and non-proliferative LN. **Supplementary Table 4** summarizes the predictive power of all univariate and specific extended logistic models, and the performance of select models is visualized in **Figure 5**. We point out that CRP has little to no discriminatory power in this setting (c statistic 0.56) and that the best univariate models are those constructed using GlycA or BMI (c statistic 0.72 and 0.75, respectively). A bivariate model using only GlycA and BMI (c statistic 0.91) outperforms a model constructed of six measures used in current clinical practice (i.e. CRP, serum albumin concentration, eGFR, proteinuria, C3 levels and evidence of anti-dsDNA, c statistic 0.90). We emphasize that GlycA and BMI independently contribute information about the inflammatory burden, evidenced by their independent significance in the multivariate model. Algorithmic construction of a logistic regression model using the exhaustive best subset approach results in a model using BMI, GlycA, presence of anti-dsDNA and daily corticosteroid dosage, capable of perfectly separating all non-proliferative from the proliferative cases.

**Figure 5:**
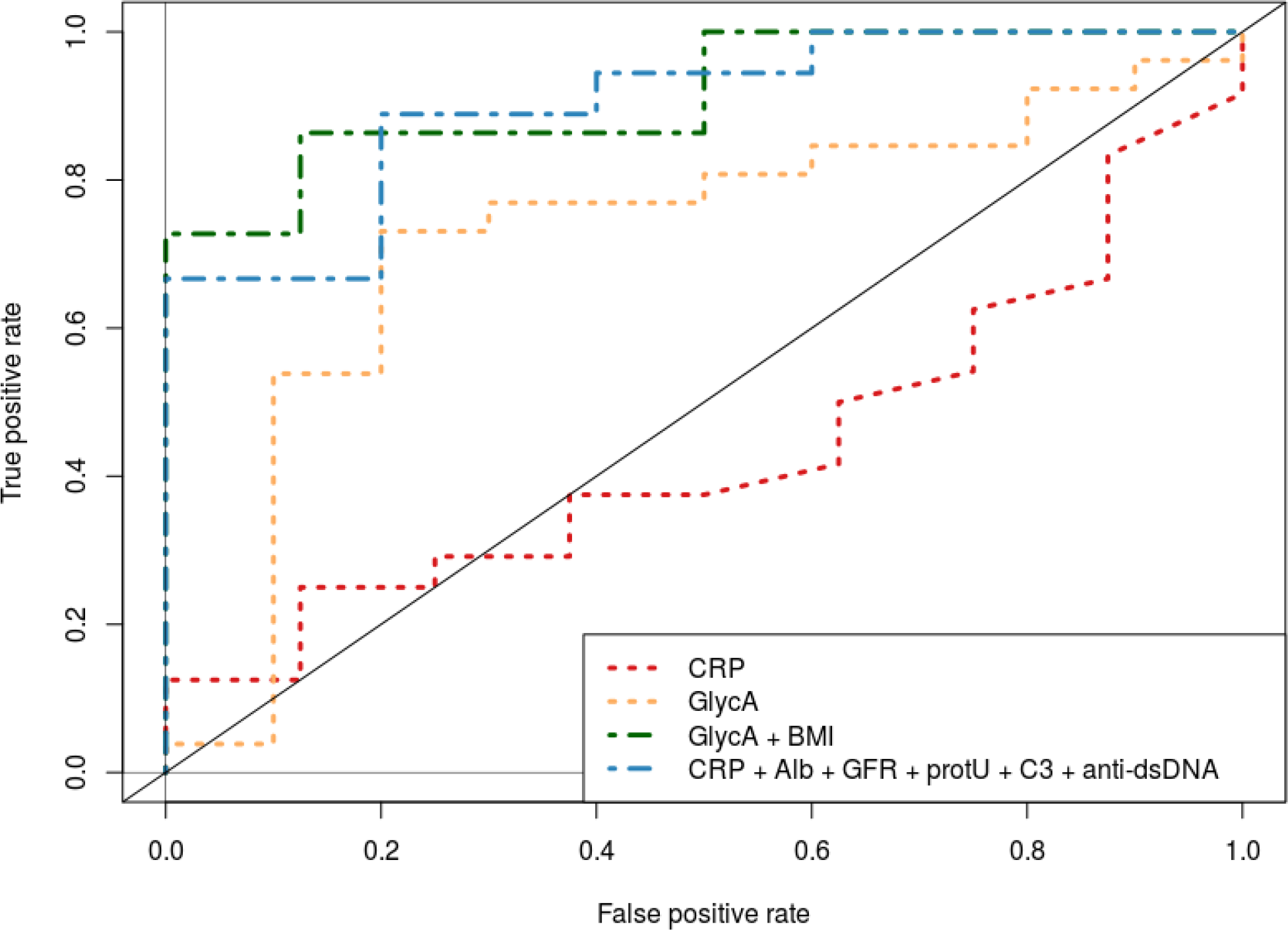
Receiver operating characteristic curve of selected logistic regression models predicting the proliferative nature of LN cases, using the variables indicated in the legend.

## Discussion

The results of the present study confirm that GlycA is elevated in SLE patients compared to healthy controls. We extend previous findings and show that GlycA level is correlated with SLEDAI and is higher in active SLE than in quiescent SLE. Moreover, in patients sampled at the time of a biopsy-proven LN flare, we show that GlycA level is higher in patients with proliferative LN than in patients with non-proliferative LN. Interestingly, although we confirm the inverse correlation between GlycA and eGFR (GlycA increasing as renal function declines), we show that the elevation of GlycA in active SLE or in proliferative LN is not solely due to an altered renal function, since non-lupus nephritic controls, who displayed lower eGFR than SLE patients, had lower GlycA levels. Moreover, the association between GlycA level and the proliferative status of LN persisted after correction for eGFR. Interestingly, after correction for other confounding factors, whether they were previously described (BMI[9], [24]–[26] and daily corticosteroid dosage[16]) or observed in this work (neutrophil count), GlycA concentration remains significantly associated with the proliferative status of flaring LN patients. The increased GlycA concentration of proliferative LN is not explained by massive proteinuria, as proteinuria levels were comparable between patients with proliferative and non-proliferative LN.

We constructed logistic regression models to determine if the GlycA marker could be of use in the prediction of the proliferative status of LN. We show that GlycA, as a robust biomarker summarizing chronic inflammatory burden from multiple sources, has high predictive value in this context. A bivariate logistic model using GlycA and BMI to predict proliferative status outperforms a model using CRP, eGFR, proteinuria, C3 levels, serum albumin concentration and the presence of anti-dsDNA.

The association between BMI and the non-proliferative status of LN in this cohort is interesting. Although obesity is not a risk factor for the development of SLE [27], the prevalence of obesity in patients with SLE is high, and higher BMI has been associated with disease activity [28], in addition to chronic inflammation and cardiovascular burden [29]. Higher BMI has also been associated with an increased risk of proteinuria in SLE patients [30], but this may either reflect active LN, or be the consequence of glomerular hyperfiltration and glomerulosclerosis induced by obesity itself [31]. Indeed, the detection of a significant proteinuria in a patient with SLE implies that a renal biopsy should be performed to differentiate a severe (proliferative) LN from a less severe (non-proliferative) form or from another kidney disease. The higher BMI observed in the group of patients with non-proliferative LN could be partly explained by the earlier development of proteinuria in obese patients, even in patients without severe active LN. The correlation of GlycA with BMI is well established [9], [24]–[26], but we emphasize that GlycA carries additional information relevant to the discrimination between proliferative and non-proliferative LN. This is demonstrated by the fact that both terms make a significant contribution to our logistic regression model and the high performance of the bivariate BMI and GlycA model.

The link between GlycA and LN severity is not altogether unexpected. We have previously shown the involvement of a set of neutrophil activation related genes with disease severity in LN [4], indicating the possible involvement of Neutrophil Extracellular Trap (NET) formation. Large scale gene correlation network analysis has shown that GlycA can also be associated to NET formation [14] and our data shows neutrophil count correlates to GlycA levels. Thus, the independent contribution of both neutrophil concentration and GlycA to the predictive logistic regression model for proliferative status is of particular interest and warrants further research into a mechanistic explanation of this phenomenon.

We point out specific limitations and strengths of this study. The limited sample sizes and unbalanced study design inherent to convenience sampling restrict the hypotheses which can be statistically tested in this dataset. Specifically, while our observations show evidence that GlycA could be an excellent candidate biomarker for treatment follow-up, the lack of rigorous periodic follow-up sampling leading up to and following the flare event obfuscate the dynamics of the GlycA marker in this period. The strengths of this study lie in the robust phenotypical characterization of patients, the completeness of clinical data recorded, and the availability of samples drawn at the time of a biopsy-proven LN flare.

The GlycA biomarker quantifies the degree to which specific acute-phase glycoproteins are acetylated. Further research is required to elucidate whether the observed GlycA associations can be ascribed to either the concentrations of these proteins, their glycosylation or their acetylation profiles. Recent evidence in cardiovascular disease (CVD) research suggests that immunoglobulin G glycosylation traits can be both positively and negatively associated to CVD risk [32], illustrating our understanding of these processes is still incomplete and inviting further study of these topics.

Our study also provides a plausible explanation of why previous studies have reported conflicting information with regards to the association between GlycA and SLE disease severity. The population studied by Chung et al [33], which reported no association to disease activity, consisted of patients with relatively low disease activity and without nephritis, while the population of Durcan et al [16], where an association to disease activity was reported, was more heterogeneous and included SLE patients with renal involvement.

Taken together, our results indicate that serum GlycA concentration, as a summary measure for multiple inflammatory processes, could be a valuable biomarker to discriminate proliferative from non-proliferative forms of lupus nephritis in flaring patients and could be used to follow-up treatment response. We believe this biomarker should be more extensively tested in large, prospective cohorts of patients with SLE. Our data suggests that GlycA could possibly be used as a globally predictive biomarker and we advocate testing its capacity to identify patients with increased risk of mortality, morbidity and disease flares over longer periods of time.

## Acknowledgement

We thank the patients who participated in the biobanks which allowed this work. We thank Sophie Tardoski, Laurent Samson and Alain Vazi, from the Centre d’Investigation Clinique of La Conception, for their help with the samples.

## Supplementary Material

**Supplementary Table 1:**
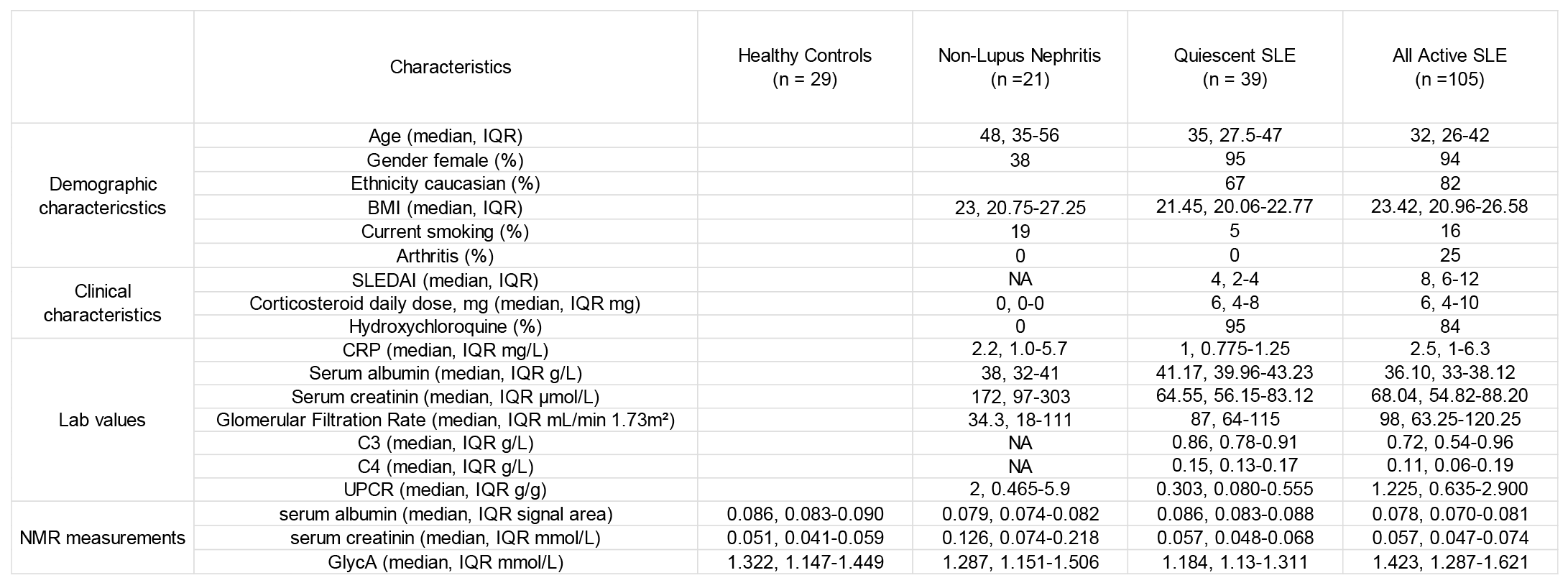
Clinical and Demographic overview

**Supplementary Table 2:**
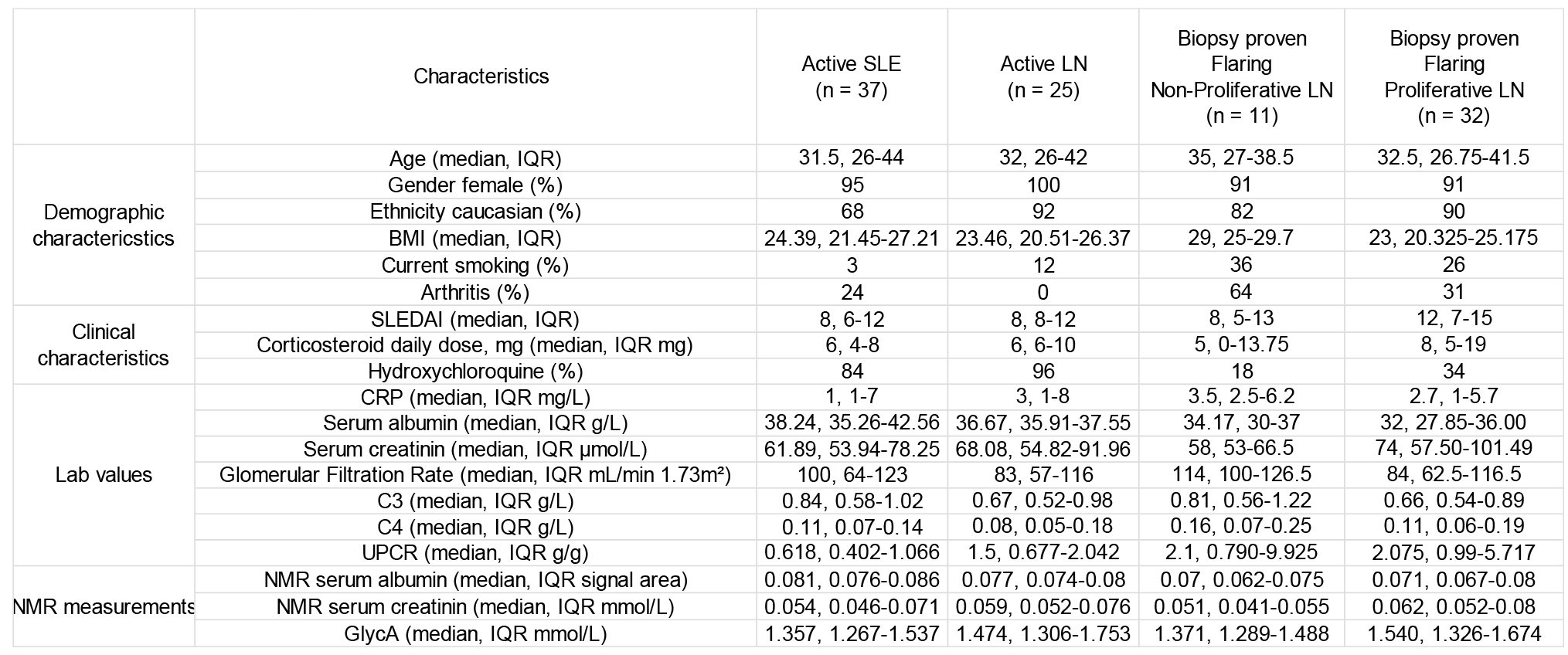
SLE patient clinical and demographical overview

**Supplementary Table 3:** Spearman correlation of GlycA to measured clinical variables. Significant adjusted p values (p<0.05 after Benjamini Hochberg multiple testing correction) are bolded.

| Characteristics | All Samples<br>N=194 |  |  | All SLE Patient<br>N=144 |  |  | Quiescent SLE<br>N=39 |  |  | Active SLE<br>N=105 |  |  | Biopsy-proven Flaring LN<br>N=43 |  |  |
| --- | --- | --- | --- | --- | --- | --- | --- | --- | --- | --- | --- | --- | --- | --- | --- |
|  | Spearman ρ | p | adjusted p | Spearman ρ | p | adjusted p | Spearman ρ | p | adjusted p | Spearman ρ | p | adjusted p | Spearman ρ | p | adjusted p |
| Albumin | -0.38 | **5.36E-06** | **1.96E-05** | -0.41 | **4.90E-06** | **1.80E-05** | -0.22 | 2.37E-01 | 5.92E-01 | -0.26 | **1.58E-02** | 6.99E-02 | 0.05 | 7.63E-01 | 7.63E-01 |
| BMI | 0.01 | 9.50E-01 | 9.50E-01 | 0.07 | 4.14E-01 | 4.45E-01 | 0.07 | 6.58E-01 | 7.18E-01 | -0.12 | 2.49E-01 | 3.42E-01 | 0.07 | 6.73E-01 | 7.63E-01 |
| C3 | -0.18 | **3.73E-02** | 5.85E-02 | -0.18 | **3.73E-02** | 8.20E-02 | -0.06 | 7.18E-01 | 7.18E-01 | -0.13 | 2.13E-01 | 3.34E-01 | -0.15 | 3.93E-01 | 7.63E-01 |
| C4 | -0.10 | 2.53E-01 | 2.79E-01 | -0.10 | 2.53E-01 | 3.10E-01 | -0.08 | 6.11E-01 | 7.18E-01 | 0.04 | 7.01E-01 | 7.71E-01 | 0.06 | 7.49E-01 | 7.63E-01 |
| CRP | 0.49 | **5.31E-10** | **5.84E-09** | 0.49 | **8.91E-09** | **9.80E-08** | 0.57 | **6.06E-04** | **6.06E-03** | 0.36 | **4.38E-04** | **4.82E-03** | 0.06 | 7.32E-01 | 7.63E-01 |
| CorticoDosage | 0.11 | 1.88E-01 | 2.30E-01 | 0.07 | 4.45E-01 | 4.45E-01 | 0.12 | 5.56E-01 | 7.18E-01 | 0.02 | 8.51E-01 | 8.51E-01 | -0.22 | 1.71E-01 | 4.71E-01 |
| Creatinine | 0.17 | **3.68E-02** | 5.85E-02 | 0.16 | 5.47E-02 | 9.83E-02 | 0.41 | **1.22E-02** | 6.09E-02 | 0.13 | 1.78E-01 | 3.26E-01 | 0.34 | **2.93E-02** | 1.61E-01 |
| GFR | -0.13 | 9.12E-02 | 1.25E-01 | -0.13 | 1.40E-01 | 1.93E-01 | -0.37 | **2.32E-02** | 7.74E-02 | -0.14 | 1.71E-01 | 3.26E-01 | -0.34 | **2.75E-02** | 1.61E-01 |
| Neutrophil | 0.34 | **1.67E-02** | **3.68E-02** | 0.35 | 6.25E-02 | 9.83E-02 | NA | NA | NA | 0.35 | 6.25E-02 | 1.72E-01 | 0.35 | 6.25E-02 | 2.29E-01 |
| SLEDAI | 0.36 | **1.20E-05** | **3.31E-05** | 0.36 | **1.20E-05** | **3.31E-05** | 0.15 | 3.53E-01 | 7.06E-01 | 0.10 | 3.38E-01 | 4.13E-01 | -0.09 | 5.65E-01 | 7.63E-01 |
| UPCR | 0.38 | **4.84E-06** | **1.96E-05** | 0.40 | **4.76E-06** | **1.80E-05** | -0.12 | 5.68E-01 | 7.18E-01 | 0.24 | **1.91E-02** | 6.99E-02 | 0.08 | 6.03E-01 | 7.63E-01 |

**Supplementary Table 4:**
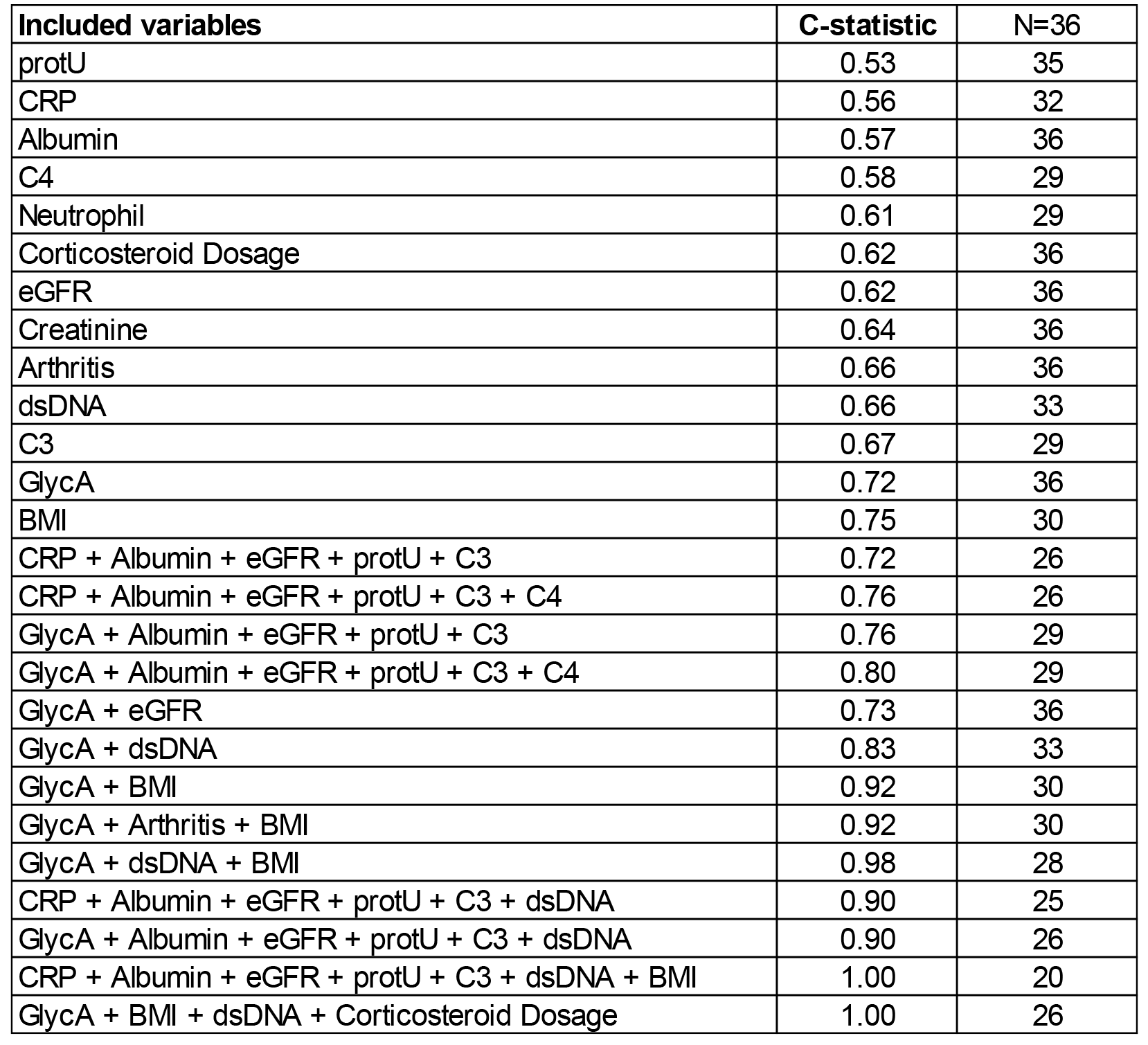
c-statistics of all univariate and selected multivariate logistic regression models

| Included variables | C-statistic | N=36 |
| --- | --- | --- |
| protU | 0.53 | 35 |
| CRP | 0.56 | 32 |
| Albumin | 0.57 | 36 |
| C4 | 0.58 | 29 |
| Neutrophil | 0.61 | 29 |
| Corticosteroid Dosage | 0.62 | 36 |
| eGFR | 0.62 | 36 |
| Creatinine | 0.64 | 36 |
| Arthritis | 0.66 | 36 |
| dsDNA | 0.66 | 33 |
| C3 | 0.67 | 29 |
| GlycA | 0.72 | 36 |
| BMI | 0.75 | 30 |
| CRP + Albumin + eGFR + protU + C3 | 0.72 | 26 |
| CRP + Albumin + eGFR + protU + C3 + C4 | 0.76 | 26 |
| GlycA + Albumin + eGFR + protU + C3 | 0.76 | 29 |
| GlycA + Albumin + eGFR + protU + C3 + C4 | 0.80 | 29 |
| GlycA + eGFR | 0.73 | 36 |
| GlycA + dsDNA | 0.83 | 33 |
| GlycA + BMI | 0.92 | 30 |
| GlycA + Arthritis + BMI | 0.92 | 30 |
| GlycA + dsDNA + BMI | 0.98 | 28 |
| CRP + Albumin + eGFR + protU + C3 + dsDNA | 0.90 | 25 |
| GlycA + Albumin + eGFR + protU + C3 + dsDNA | 0.90 | 26 |
| CRP + Albumin + eGFR + protU + C3 + dsDNA + BMI | 1.00 | 20 |
| GlycA + BMI + dsDNA + Corticosteroid Dosage | 1.00 | 26 |

**Supplementary Figure 1:**
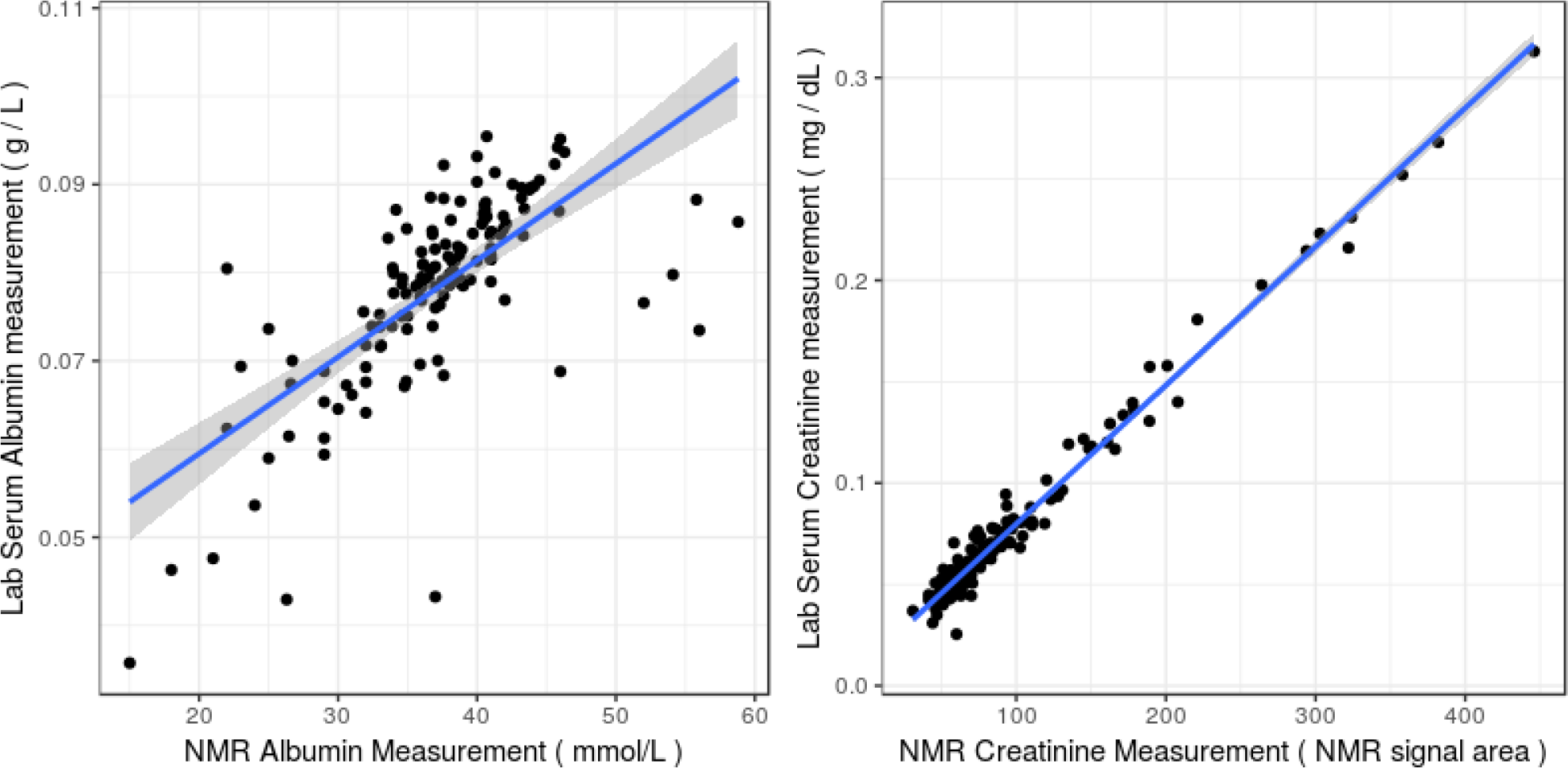
Measurements of serum creatinine and albumin levels correlate very well between NMR and standard laboratory tests (*ρ*=0.74 and *ρ*=0.94, respectively, with p-values<10^−8^).

